# Understanding the phytochemical diversity of plants: Quantification, variation and ecological function

**DOI:** 10.1101/2023.03.23.533415

**Authors:** Hampus Petrén, Redouan Adam Anaia, Kruthika Sen Aragam, Andrea Bräutigam, Silvia Eckert, Robin Heinen, Ruth Jakobs, Lina Ojeda-Prieto, Moritz Popp, Rohit Sasidharan, Jörg-Peter Schnitzler, Anke Steppuhn, Frans Thon, Sebastian Tschikin, Sybille B. Unsicker, Nicole M. van Dam, Wolfgang W. Weisser, Meike J. Wittmann, Sol Yepes, Dominik Ziaja, Caroline Müller, Robert R. Junker

## Abstract

Plants produce a great number of phytochemical compounds mediating a variety of different functions. Recently, phytochemical diversity (chemodiversity), a way which to quantify the complex phenotype formed by sets of phytochemicals, has been suggested to be important for function. However, no study has systematically examined the potential (in)direct functional importance of chemodiversity on a general level, partly due to a lack of an agreement on how to quantify this aspect of the plant phenotype. This paper has four aims: 1) We discuss how chemodiversity (deconstructed into components of richness, evenness and disparity) may quantify different aspects of the phenotype that are ecologically relevant. 2) We systematically review the literature on chemodiversity to examine methodological practices, explore ecological patterns of variability in diversity across different levels of biological organization, and investigate the functional role of this diversity in interactions between plants and other organisms. 3) We provide a framework facilitating decisions on which measure of chemodiversity is best used in different contexts. 4) We outline open questions and avenues for future research in this area. A more thorough understanding of phytochemical diversity will increase our knowledge on the functional role phytochemical compounds, and how they shape ecological interactions between plants and their environment.

## Introduction

The phytochemical compounds produced by plants are crucial for shaping interactions between plants and their surrounding environment (Fraenkel 1959, Hartmann 2007). Individual compounds can be seen as functional traits, that impact the physiology, interactions and ultimately fitness of plants (Müller and Junker 2022, Walker et al. 2022). Together, a mixture of phytochemicals form a complex phenotype that may vary along multiple dimensions (Marion et al. 2015). These dimensions include the total number of phytochemical compounds included in a mixture, the quantitative variation in the abundance of those compounds, and the biosynthetic origins and molecular structures of them. Traditionally, most of the attention around the function of phytochemicals has been focused on the effect of individual compounds on the interaction between a plant and a specific interacting organism (Richards et al. 2010). More recently, studies aiming to more comprehensively measure the phytochemical phenotype have found that other aspects of it, such as the number and relative abundances of compounds, are important for different functions, and ultimately plant fitness (Junker et al. 2018, Dyer et al. 2018). However, despite, or perhaps due to, the complex nature of the phytochemical phenotype, we lack a more comprehensive understanding of what dimensions of it are important for different functions in different types of ecological interactions.

A method of characterizing the phytochemical phenotype that is increasingly often used is to measure its diversity (Wetzel and Whitehead 2020). By utilizing diversity indices, a multivariate phenotype can be summarized into a univariate measure of its diversity, which quantifies a certain aspect of the phenotype (Marion et al. 2015, Petrén et al. 2023). Such a measure of phytochemical diversity (chemodiversity; see Box 1 for a glossary of central terms) can be useful if it encompasses biologically meaningful variation that is associated with the phenotype’s function. In this article, we review phytochemical diversity, how it has been measured in published studies, and provide recommendations for how to analyse it. Doing so, we highlight ways of quantifying the phytochemical phenotype that are ecologically relevant, suitable to different types of data and suited to answer specific research questions.

### The phytochemical phenotype

At least 200 000 phytochemical compounds have been described, and many more are sure to exist (Kessler and Kalske 2018, Wang et al. 2019). Depending on the species of plant, what part of the plant is examined, and the method by which compounds are extracted and identified, anything from just a few to several thousand compounds may be found in a sample (Uthe et al. 2021, Li and Gaquerel 2021). An individual phytochemical compound will occur at a particular abundance, originate from a specific biosynthetic pathway, and have a certain molecular structure. Regardless of the context, phytochemicals occur in mixtures, with variation in the total number and abundances of compounds. Therefore, suitable methods of quantifying and summarizing this multivariate phenotype are needed. These measures of the phenotype should ideally be linked to its function, such that they are associated with plant performance and fitness. Figuring out what dimensions of the phenotype are most important in this regard will increase our understanding of how phytochemicals shape interactions between plants and their environment.

There are many examples where different aspects of the phytochemical phenotype are important for function. Often, studies focus on a univariate dimension of the phenotype, and associate this with function. Most commonly, this is the presence or abundance of specific compounds. For example, single compounds have been shown to limit bacterial growth, reduce herbivory or increase pollination success (Lankau 2007, Zhou et al. 2017, Burdon et al. 2018). In other cases, function is linked to the total abundance of compounds, or derive from combinations of compounds (Duffey and Stout 1996, Gershenzon et al. 2012, Calf et al. 2018). For example, a combination of several compounds may be necessary for optimal herbivore defence or pollinator attraction (Berenbaum et al. 1991, Byers et al. 2014). Other studies have linked function with principal components of the mixture of phytochemicals (Poelman et al. 2009), or found that function may depend on compounds occurring in specific ratios (Berenbaum and Neal 1985, Junker et al. 2018, Orlando et al. 2022). In the past decade, interest in summarizing the phytochemical phenotype by using measures of chemodiversity has increased (Moore et al. 2014, Hilker 2014, Dyer et al. 2014, Marion et al. 2015, Kessler and Kalske 2018, Wetzel and Whitehead 2020, Müller et al. 2020). In this way, diversity, measured for complete sets or selected classes of phytochemicals found in plants, most often in leaves, has been linked to ecological function in a wide range of studies. For example, a higher diversity of phytoalexins was associated with a lower risk of fungal infection in in *Phaseolus* seedlings (Lindig-Cisneros et al. 2002). Combined, the described studies indicate that what aspects of the phytochemical phenotype are associated with function, and how that influences ecological interactions, can vary substantially. If the “wrong” aspect of the phenotype is measured in a given context (i.e. one that is not important for function), this might contribute to results suggesting that secondary metabolites overall have little to no effect on herbivore susceptibility (Carmona et al. 2011). Overall, it is increasingly evident that the phytochemical diversity might be a key part of the phenotype in many cases, as suggested by a rapidly increasing amount of research on the topic (Table S1). However, this diversity can be measured in different ways, and consists of different components that may be more or less ecologically relevant. Most studies have not considered such issues when quantifying the diversity of phytochemical phenotypes, which might impede efforts of linking phytochemical variation to biological function. Therefore, a closer examination of what different measures of diversity actually measure, and which indices are useful to answer different questions in different contexts is needed. In the following sections, we (1) contextualize and compare methods of quantifying the diversity of the phytochemical phenotype; (2) systematically review the literature on phytochemical diversity, elucidating general patterns on the importance of it for different ecological interactions; (3) provide recommendations for how to optimally measure this diversity for different ecological contexts, types of data and research questions; and (4) examine avenues for future research on the subject.

### Use of diversity indices in chemical ecology

Diversity indices have a long tradition of use in community ecology, where they are often used to quantify the diversity of species, which is subsequently associated with community and ecosystem level processes (Magurran 2004; Box 2). More recently, the same statistical measures have become more widely applied in biology, and have been used to quantify the diversity of e.g. elements, molecules, genes, transcriptomes, phenotypes and soundscapes (Martínez and Reyes-Valdés 2008, Kellerman et al. 2014, Marion et al. 2015, Sherwin et al. 2017, Fernández-Martínez 2022, Luypaert et al. 2022). In the field of chemical ecology, measuring chemical diversity is becoming increasingly common. However, with a few exceptions (Wetzel and Whitehead 2020, Bakhtiari et al. 2021, Ramos et al. 2023, Petrén et al. 2023), little attention has been paid to how phytochemical diversity is actually measured, and how this is relevant to its function in different ecological contexts.

What diversity is and how to best measure it is a topic of much discussion (Jost 2006, Tuomisto 2010, Morris et al. 2014, Chao et al. 2014). In this review, we focus on measures of α-diversity (“local diversity”), which in the case of chemodiversity is the diversity of a single sampling unit, most often an individual plant. This contrasts to γ-diversity (“regional diversity”) which is the total diversity at the scale of a group of sampling units, and β-diversity, which is derived from the other two measures and represents the variation, turnover or dissimilarity between sampling units (Ellison 2010, Anderson et al. 2011). On a fundamental level, diversity can be deconstructed into three components: richness, evenness and disparity (Purvis and Hector 2000, Daly et al. 2018) (Fig. 1a). For measures of chemical diversity, richness is simply the number of compounds found in a sample. The second component of diversity, evenness, is a function of relative abundances. For a set of compounds, evenness is maximised when they are all equally abundant, and decreases as abundances become more uneven. The third component, disparity, describes how different the measured entities are to each other regarding defined properties. For chemical compounds, disparity, which we will refer to as compound dissimilarity, may be based on some kind of property of the compounds that is considered relevant, such as their molecular structure or the biosynthetic pathways they were produced in (Sedio 2017, Whitehead et al. 2021, Petrén et al. 2023). In practice, this component has only rarely been measured.

**Figure 1.**
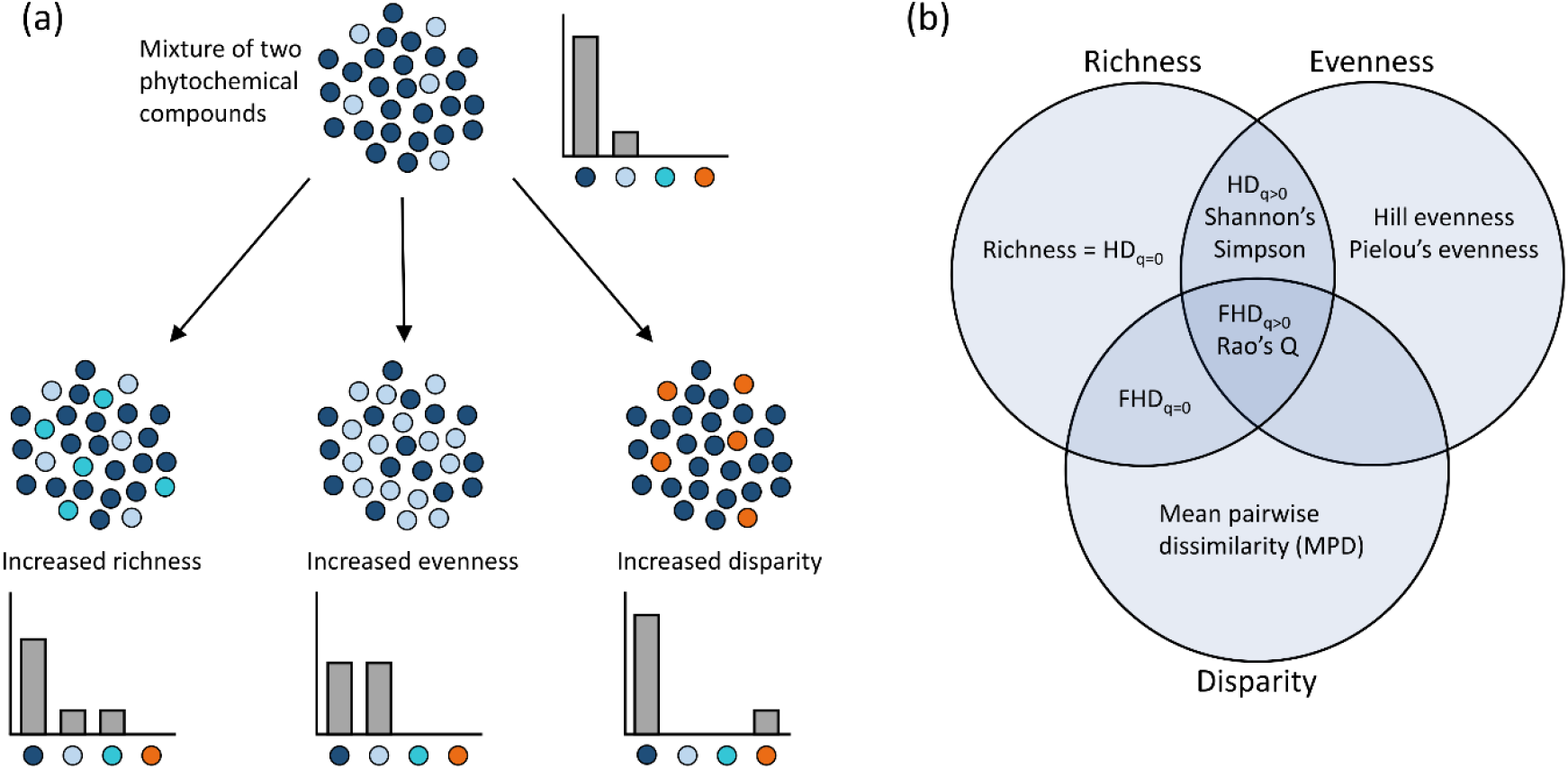
An illustration of (a) different components of diversity, and (b) some indices used to measure diversity. (a) Diversity can be deconstructed into components of richness, evenness and disparity. Here, phytochemical compounds are illustrated as points of different colours. Bar plots next to clouds of points indicate the relative abundance of each type. The top cloud of points has two different compounds with an uneven distribution. With increased richness, the number of different phytochemicals become three instead of two. With increased evenness, the relative abundances of the phytochemicals become more equal. With increased disparity, the phytochemicals become more dissimilar, e.g. in regards to molecular structure or biosynthesis (here represented as more dissimilar colours). (b) Diversity can be measured by quantifying its components, or combinations thereof. Each circle represents one component of diversity. Indices in overlapping areas measure combinations of these components. There exists a multitude of diversity indices, and we have only included ones that are considered in the main text. Abbreviations: HD_q=0_: Hill diversity at *q = 0*, which is equal to the richness; HD_q>0_: Hill diversity at *q > 0*, which is a function of richness and evenness; FHD_q=0_: functional Hill diversity at *q = 0*, which is a function of richness and disparity; FHD_q>0_: functional Hill diversity at *q > 0*, which is a function of all three components of diversity.

Many indices can be used to quantify diversity, each with unique properties and mathematical behaviours that may emphasize different components of it (Fig. 1b). For phytochemical diversity, to simply quantify it as richness is the most basic measure, and is frequently used (Table S1). Direct quantifications of evenness are more rare. The most common way of quantifying chemodiversity is with an index that combines richness and evenness. Using the relative abundances of the compounds in a sample, indices such as the Shannon’s diversity index or the inverse Simpson diversity can be calculated. Functional diversity indices, which include the disparity component of diversity (in the form of a dissimilarity matrix), have only been used in a few cases. This includes measures of mean pairwise dissimilarity (MPD; the mean of values in the compound dissimilarity matrix), which only include the disparity component (Whitehead et al. 2021, Volf et al. 2022), and indices such as Rao’s Q that are dependent on all three components of diversity (Bakhtiari et al. 2021). A more detailed mathematical description of these indices in the context of phytochemical diversity is available in Petrén et al. (2023).

An underappreciated problem in many studies is that different diversity indices measure different things (Wetzel and Whitehead 2020). An index necessarily emphasizes some components of diversity while de-emphasizing others, which may affect interpretations of the results (Tuomisto 2010, Steel et al. 2013). For example, Bakhtiari et al. (2021) found that only some measures of chemical diversity of glucosinolates in *Cardamine* species were associated with herbivore performance. This, they argue, indicates that different aspects of the diversity might have different ecological effects. Most studies however calculate only a single index, often without justifying the choice. This is not ideal, since a certain index might emphasize an aspect of the phenotype that happen to be less ecologically relevant. Instead, it might be better to calculate diversity using measures where it is clear which component(s) of diversity is actually quantified.

In community ecology, many agree that diversity is optimally measured using Hill diversity (Ellison 2010). Hill diversity, also called Hill numbers or effective numbers, is closely related to traditional diversity indices such as Shannon’s diversity, but offer several advantages (Hill 1973, Jost 2006, Chao et al. 2014). This includes intuitive units, easy partitioning into α-, β- and γ-diversity, and options to quantify functional diversity (Chiu and Chao 2014, Chao et al. 2014). Additionally, Hill diversity includes a parameter, *q*, called the diversity order, which controls how sensitive the measure is to the relative abundances of compounds. By selecting the type of Hill diversity to calculate and the value of *q*, a researcher can calculate a measure of diversity that focuses on any combination of its three components (richness, evenness and disparity). This allows for both easier and more extensive analyses of diversity, compared to using just a single traditional index.

While chemical diversity has primarily been measured using traditional indices, Hill diversity has been increasingly used in recent years. Marion et al. (2015) introduced the concept to characterize phenotypic complexity. Other studies followed in measuring phytochemical diversity in this way (Harrison et al. 2016, 2018, Glassmire et al. 2016, Cosmo et al. 2021, Philbin et al. 2021, 2022). Functional Hill diversity, which includes compound dissimilarities in diversity calculations, has only very recently been used to quantify chemical diversity. Forrister et al. (2022) measured the diversity of metabolites in leaves of *Inga* species, using compound dissimilarities based on cosine similarities between MS/MS spectra. Petrén et al. (2023) developed the R-package *chemodiv*, which provides functions to aid chemical ecologists to more comprehensively quantify chemical diversity for a wide range of datasets. This includes functional Hill diversity, where compound dissimilarities, calculated based on molecular and/or biosynthetic properties of the compounds, are included in diversity calculations. Overall, as phytochemical diversity is increasingly often quantified, there is a need to closer examine what components of this diversity are most ecologically relevant, and consider the mechanisms by which diversity is important for function (Wetzel and Whitehead 2020, Bakhtiari et al. 2021).

### Systematic literature review

To compile information on the use of diversity indices in quantifying phytochemical diversity, we conducted a systematic search of literature on the subject. In February 2023, we searched the Web of Science for studies on phytochemical diversity with separate searches of *“*chemical diversity” and plant\**, *chemodiversity and plant\**, *“secondary compound” or “secondary metabolite” and diversity and plant\**, and *“voc diversity” or “scent diversity” or “volatile diversity” and plant\**. These searches returned a total of 2106 scientific articles. These were screened by title and abstract, and potentially relevant papers were examined in detail for use of diversity indices quantifying mixtures of phytochemical compounds. In addition to the literature search, we screened all papers cited by, and all papers that cited a number of key papers in the field (Moore et al. 2014, Hilker 2014, Marion et al. 2015, Kessler and Kalske 2018, Wetzel and Whitehead 2020, Whitehead et al. 2021, Bakhtiari et al. 2021, Cosmo et al. 2021, Philbin et al. 2022), and included a small number of additional papers known to the authors but not detected in the systematic search. We included studies quantifying α-diversity, or some component (richness, evenness, disparity) of it. Most often, this represents the chemodiversity of individual plants, and is therefore a measure of the phenotype, but in some cases this includes lower (within plant) or higher (population or community) level variation, and such studies were included if this was considered as the sampling unit by the authors. Studies quantifying only β-diversity or dissimilarity were not included. Although we believe our search to be exhaustive in regards to finding studies using diversity indices, it should be noted that many studies reporting only phytochemical richness were likely not included, as richness is often not framed in the context of diversity, and instead only briefly reported in the results. We only included studies quantifying richness if they did so in the context of diversity. Our collection of studies includes both those examining variation in phytochemical diversity (comparing populations, species, treatments etc.) and studies examining function through the effect of phytochemical diversity on an interacting organism (e.g. effects on herbivore performance or fungal growth). In total, we found 87 studies in our systematic search that fitted the criteria. Table S1 lists these studies, including the study system, type of phytochemicals measured, analytical method, type of measure calculated, and if variation in or an effect of the diversity was found. Below, we consider theoretical aspects of each component of diversity separately and combined, and review the empirical studies that have measured variation in and the function of this diversity, synthesising patterns of variation and potential mechanisms of function.

### Phytochemical richness

Richness, the number of compounds, is the most straightforward component of chemodiversity. The number of compounds a plant species produces should be largely dependent on the number of enzymes that make up the corresponding biosynthetic pathways. Therefore, from an evolutionary perspective, phytochemical richness may represent an “evolutionary potential”, where a high phytochemical diversity could reflect and ability for adaptive responses to selection in the same way as genetic diversity. On the level of individual plants, a high phytochemical richness may also enable higher levels of phenotypic plasticity in response to changing conditions.

From an ecological perspective, producing a high number of phytochemicals might benefit a plant through mechanisms which are linked to major hypotheses aiming to explain how phytochemical diversity is maintained by natural selection (Wetzel and Whitehead 2020, Whitehead et al. 2021). According to the “interaction diversity hypothesis”, for a plant experiencing a multitude of interactions, e.g. being attacked by multiple herbivores, producing more compounds is useful if different compounds are efficient against different herbivores. This would then result in selection for increased phytochemical richness (Berenbaum and Zangerl 1996, Iason et al. 2011, Junker 2016). If, as suggested by the “screening hypothesis”, most compounds are instead non-functional, a high richness may increase the probability that at least some compounds in a mixture are functional (Jones and Firn 1991, Firn and Jones 2003, but see Pichersky et al. 2006). These two scenarios suggest that producing a higher number of compounds may benefit plants experiencing multiple interactions. In these cases, function may still originate from the effect of individual compounds. However, under the “synergy hypothesis” phytochemical richness *per se* may be mechanistically important in single interactions. Synergistic effects, where the effect of a mixture of compounds is greater than the combined effects of individual compounds, have been documented in a number of systems, and may emerge via several different mechanisms (Richards et al. 2016). The probability of synergistic effects occurring may increase with the number of compounds present in a mixture, thereby creating selection for increased phytochemical richness.

The main advantage of quantifying chemodiversity as compound richness is that, compared to other measures, it is straightforward to measure. There are a number of disadvantages though. First, levels of measured richness may vary widely depending on what methods are used for extraction and detection of compounds (Uthe et al. 2021), and increase with the total abundance of phytochemicals in a plant, for both technical and biological reasons (Wetzel and Whitehead 2020). Overall, the measured richness will be an underestimation of the true metabolic space of a plant (Li and Gaquerel 2021). This restricts comparisons of phytochemical richness to cases where similar analytical methods have been used. Second, measures of richness disregard the relative abundances of compounds. This means that compounds occurring at low abundances, which may be functionally less important in some contexts, contribute as much to the measure of richness as high abundance compounds.

Phytochemical richness is a frequently reported measure. Variation in it has been documented on all levels of biological organization, including between tissues (Whitehead et al. 2013, Elser et al. 2022), individuals (Ziaja and Müller 2022), populations (Zeng et al. 2022, Eisen et al. 2022), species (Macel et al. 2014, Züst et al. 2020), orders (Courtois et al. 2009), and communities (Peguero et al. 2021), as well as across herbivory treatments (Agrawal 2000), phylogenies (Becerra et al. 2009, Cacho et al. 2015) and landscapes (Defossez et al. 2021). Here, richness varies from a handful of compounds of a specific biosynthetic class, to several thousands metabolic features that are assumed to represent individual unidentified compounds.

Whereas some studies solely document variation in phytochemical richness, others go a step further and investigate the functional importance of this variation. For example, a higher phytochemical richness has been shown to reduce preference and/or performance of specific herbivores in a number of study systems (Adams and Bernays 1978, Castellanos and Espinosa-García 1997, Agrawal 2000). In natural environments, phytochemical richness, both on the level of individual plants and whole communities, has been found to shape ecological interactions, with effects such as reducing species richness of herbivores (Salazar et al. 2018), decreasing arthropod abundances (Defossez et al. 2021) and reducing levels of herbivore damage (Whitehead et al. 2013). This suggests that phytochemical richness may be important for function in both specific interactions, through synergistic effects between compounds, and that it may increase herbivore resistance of plants experiencing pressures from multiple herbivores, through different compounds being efficient against different herbivores.

### Phytochemical evenness

Evenness is dependent on the relative abundances of compounds. Compared to richness, direct measures of evenness are rare. It is conceivable that evenness is a comparatively more plastic component of the phytochemical phenotype, as up- or down-regulation of specific biosynthetic pathways may be more likely to affect the relative abundances of compounds than the total number of compounds. Mechanistically, evenness might be important for function in different ways. If function is dependent on synergies between compounds occurring in roughly equal abundances, a set of compounds with high evenness might enable more/stronger synergies than a set of compounds with low evenness. On the other hand, if function comes from a few specific compounds, a high evenness may be disadvantageous if it reflects a lower abundance of those compounds (Pais et al. 2018, Wetzel and Whitehead 2020).

Although a potentially interesting measure, evenness has a number of disadvantages. First, accurately quantifying the relative abundances of structurally different molecules in a chromatogram can be challenging (Walker et al. 2022). Additionally, calculations of relative abundances could be based on the mass concentrations, or on molar concentrations, which may result in different relative abundances of compounds in phytochemical mixtures where molecules vary extensively in molecular mass. Second, if the bioactivity of compounds varies widely, such that also low-abundance compounds are important for function, measuring their evenness is less relevant (Clavijo McCormick et al. 2014). Third, despite seemingly a straightforward measure, there is no consensus on how to best quantify evenness (Smith and Wilson 1996, Jost 2010, Tuomisto 2012, Chao and Ricotta 2019). Many different measures of it exist, with different mathematical properties (Smith and Wilson 1996). The most popular measure is likely Pielou’s evenness (Pielou 1966), but evenness can also be calculated in the Hill diversity framework (Hill 1973). The behaviour of these two indices can differ, with the former measure not being truly independent of richness (DeBenedictis 1973, Alatalo 1981), and there is no agreement on which version is most suitable (Jost 2010, Tuomisto 2012).

Among the few studies that have quantified phytochemical evenness, there is evidence of variation in evenness of (classes of) compounds between bryophyte species (Peters et al. 2019, 2021), differences in cardenolide evenness between wild type and mutant *Erysimum cheiranthoides* (Mirzaei et al. 2020), and differences between maize plant types (Bernal et al. 2023). Pais et al. (2018) found that an increased evenness of leaf metabolites in *Cornus florida* was associated with a higher probability of plants being diseased. In contrast, Feng et al. (2021) noted a positive association between evenness and antibacterial activity for *Juniperus rigida* essential oils, and Whitehead et al. (2021) found that increasing the evenness of phenolics in the diet of different insect herbivores increased how many of them that were negatively affected by the phenolics. Measuring covariation in the diversity of plants, arbuscular mycorrhizal fungi and arthropods, as well as chemical and genetic diversity associated with *Plantago lanceolata* individuals, Morris et al. (2014) found that evenness showed different patterns compared to richness and e.g. Shannon’s diversity, suggesting it represents different information not captured by other diversity indices.

Overall, the limited evidence from studies so far suggests that effects of phytochemical evenness might differ between study systems. As basically any kind of variation in the phytochemical phenotype will affect evenness, more research is needed to investigate potential mechanisms and its importance for ecological interactions in different contexts. Additionally, in the studies cited above, four different indices of evenness were used. As a consequence, effects of evenness could differ not only depending on ecological context, but also due to what index is used. This illustrates the challenges of measuring this component of the phytochemical phenotype, and in interpreting results (Wetzel and Whitehead 2020).

### Phytochemical disparity

Disparity (compound dissimilarity) is rarely quantified in studies of chemodiversity, but may often be an important component of it. Dissimilarities between chemical compounds may be calculated in a number of different ways, based on e.g. their molecular substructures (Cao et al. 2008), physicochemical properties (Dowell and Mason 2020), molecular fingerprints (Cereto-Massagué et al. 2015) or what biosynthetic pathways or enzymes produce them (Junker 2018, Petrén et al. 2023). Additionally, recently developed methods in the GNPS ecosystem (Wang et al. 2016) enable calculations of compound dissimilarities for unidentified compounds. Here, cosine dissimilarities are calculated based on comparisons of mass spectra (Wang et al. 2016, Aksenov et al. 2021), a method which has been applied in chemical ecology as well (Sedio 2017). In an ecological context, a crucial assumption for why compound dissimilarities are relevant is that there is an association between the structure of a compound and its function. Generally, molecules with a similar structure can be expected to on average have a more similar biological activity/function compared to molecules with a more different structure (Berenbaum and Zangerl 1996, Martin et al. 2002). Additionally, more dissimilar compounds may be more likely to function synergistically than less dissimilar compounds (Liu and Zhao 2016). Therefore, a structurally diverse set of phytochemicals may be more functionally diverse (e.g. efficient against a larger set of herbivores) and/or potent (e.g. efficient at lower abundances) (Philbin et al. 2022). From an evolutionary perspective, instead considering compound dissimilarities based on biosynthetic pathways may be useful (although structural and biosynthetic similarity are often correlated). In this case, if a plant species produces a set of compounds with a high average compound dissimilarity, this indicates that these are produced in multiple biosynthetic pathways, indicating that the species has an extensive metabolic machinery for producing phytochemicals. Comparing such compound dissimilarities across phylogenies may generate insights on the evolution of phytochemicals.

We believe that including compound dissimilarities in measures of chemodiversity can be meaningful. However, there are number of challenges associated with it. First, as mentioned, its ecological usefulness rests on the assumption of a link between structure and function, such that two compounds that are structurally dissimilar are also functionally dissimilar. Although this may not the case for all pairs of phytochemicals in a set of compounds, an association between the structural diversity (level of disparity) for the whole set of compounds and its diversity in function may be more likely. Second, related to the first point, compound dissimilarity can be quantified in many different ways. We have made a distinction between compound dissimilarities calculated based on biosynthesis and molecular structure, but there are many alternative ways of quantifying compound dissimilarities based on their structures (Cao et al. 2008, Cereto-Massagué et al. 2015). Overall, more research is needed to examine the link between (different measures of) the structure and function of phytochemical compounds.

We are aware of two studies that have specifically quantified the disparity/compound dissimilarity of sets of phytochemicals as a measure of their structural diversity. Similar to results on evenness, Whitehead et al. (2021) found that an increased structural diversity of phenolics, quantified as the mean pairwise dissimilarity (MPD) of all compounds in a mixture, increased the proportion of consumers negatively affected by the compounds. Studying different *Salix* species along elevational gradients, Volf et al. (2022) found an increase in the structural diversity (MPD) of salicinoids with altitude, but a decrease for flavonoids, which may result from variation in abiotic factors. Although evidence is limited so far, these studies suggest that structural diversity can be an important aspect of the phytochemical phenotype that varies with environmental conditions and affects function.

In addition to using compound dissimilarities to examine structural diversity, it should be noted that other studies have quantified dissimilarities or properties of phytochemicals in other contexts. Sternberg et al. (2012) and Rasmann (2014) examined effects of the polarity of phytochemicals on herbivore resistance, with evidence indicating that apolar cardenolides may be more toxic to herbivores than polar cardenolides. Cosine dissimilarities between MS/MS spectra of compounds have also been used in a number of studies to calculate compound dissimilarities. These have then been included in calculations of measures that integrate structural and compositional dissimilarities of sets of phytochemicals, in order to quantify such differences within and between species at different scales (Sedio et al. 2017, 2018, 2020, 2021, Ernst et al. 2019, Endara et al. 2022, Forrister et al. 2022). Although these examples regard dissimilarity between rather than diversity within sets of phytochemical compounds, they similarly point to the importance of structural variation of phytochemicals at different levels. Overall, this signals the need for more in-depth examinations of the structural component of chemodiversity and its link to ecological function.

### Phytochemical diversity – measures of combined components

While each of the three components of diversity may be measured separately, diversity indices combine the components. Shannon’s diversity, Simpson diversity or Hill diversity, which combine richness and evenness into a single measure, are often used to quantify chemical diversity. Richness and disparity may be measured in combination as the sum of all the pairwise dissimilarities in the dissimilarity matrix (Walker et al. 1999). Richness, evenness and disparity can be combined with the use of functional diversity indices, such as Rao’s Q (Rao 1982) or functional Hill diversity (Chiu and Chao 2014, Chao et al. 2014).

By combining multiple components of diversity, such indices are advantageous in that they summarize different aspects of the phytochemical phenotype in a single measure, which can be associated with function. This can be especially useful if the function of a mixture of phytochemicals is dependent on a combination of high richness, evenness and/or disparity. For example, a higher number of compounds might increase herbivore resistance for a plant, but only if the compounds occur at similar abundances, and/or are structurally dissimilar. In this case, richness alone may not correlate with herbivore resistance, while an index that includes also evenness and/or disparity will. There are also potential disadvantages with using indices. By combining multiple components of diversity, they may conceal independent variation in single components. Additionally the relative weights of different components in contributing to an index can vary. For example, Simpson diversity puts less weight on low-abundance compounds than Shannon’s diversity, and a choice has to be made about which index is most appropriate. This is made easier by the use of Hill diversity, where the selection of the diversity order (*q*) controls the sensitivity of the index to the relative abundances of compounds. More than anything, it is important to understand the properties of the indices calculated, as that will help understanding how different aspects of the phytochemical phenotype mechanistically contributes to function.

Studies quantifying chemodiversity using a diversity index most often calculate diversity as a function of richness and evenness, by calculating the Shannon’s or Simpson diversity. Similar to richness, variation in diversity has been documented at different levels of biological organization, including for different plant tissues (Eilers 2021, Elser et al. 2022), populations (Bravo-Monzón et al. 2014, 2018) and species (Ortiz et al. 2019, Peguero et al. 2021). There is also variation along altitudinal gradients (Glassmire et al. 2016, Volf et al. 2020, Philbin et al. 2021) and for different levels/types of herbivory (Li et al. 2020, Philbin et al. 2022). Among studies that have examined functional effects, results indicate that an increased phytochemical diversity can decrease levels of herbivory (Richards et al. 2015, Glassmire et al. 2019), decrease herbivore performance (Tewes et al. 2018, Whitehead and Poveda 2019) and increase resistance to fungi and bacteria (Lindig-Cisneros et al. 1997, 2002, De-la-Cruz-Chacón et al. 2019, Feng et al. 2021). It can also affect the diversity or structure of the community of herbivores feeding on host plants (Richards et al. 2015, Volf et al. 2018, Harrison et al. 2018, Cosmo et al. 2021), affect tri-trophic interactions (Wan et al. 2017, Slinn et al. 2018), and shape the diversity of the surrounding plant and microbe communities (Iason et al. 2005, Zhang et al. 2023). Other studies have instead found that intermediate or high levels of phytochemical diversity might be disadvantageous (Sternberg et al. 2012, Pais et al. 2018), found no/limited effects of it on herbivory (Torres-Gurrola et al. 2011, Schuldt et al. 2012, Espinosa-García et al. 2021), or found no differences between different populations (Wolf et al. 2012). On balance, results from these studies suggest that an increased phytochemical diversity is often, but not always, beneficial for plants. Given that the number of studies on this topic is still limited, and that the ones that exist are done in a very diverse set of contexts and have measured different ecological effects, it is difficult to make any general conclusions about factors that may affect whether phytochemical diversity is beneficial or not.

Only a few studies have measured chemodiversity with indices that directly include also the disparity component. Bakhtiari et al. (2021) measured phytochemical diversity of glucosinolates in *Cardamine* plants with the Rao’s Q index, where compound dissimilarities were quantified based on chemical classes and molecular weights. They found variation in this measure of diversity among groups of *Cardamine* species, but no association between it and the level of resistance to different herbivores. Investigating leaf metabolites from close to 100 *Inga* species, Forrister et al. (2022) measured functional Hill diversity and compared it to a null model, concluding that plants invested in producing structurally diverse sets of compounds. It should also be noted that some of the studies cited above (e.g. Richards et al. 2015, Cosmo et al. 2021, Philbin et al. 2022) have measured phytochemical diversity in a way that (indirectly) takes compound structure into account by calculating diversity from ^1^H-NMR spectra. This method has the advantage that the measures partly depend on both the intra- and inter-molecular complexity of compounds in a sample, but the disadvantage that the different components of diversity cannot be easily separated. Overall, although the number of studies so far is limited, there is some evidence that the disparity component, in these cases based on compound structure, is an important part of chemodiversity that should be included in studies of it.

### Recommendations for measuring phytochemical diversity in different contexts

In the previous section, we reviewed the different components and indices of diversity, and compiled studies that have examined variation and ecological effects of this diversity. From this, it is clear that there is a wide variation in how phytochemical diversity is quantified. However, there is often a lack of justification as to why diversity is measured in a certain way in a certain study. As diversity consists of different components that may or may not be included in different measures of it, which may or may not be ecologically relevant, this limits our understanding of what aspects of the phytochemical phenotype are most important for function. Although how to best measure diversity is a complex issue, we believe there are some general practices that should be used. Therefore, we provide a number of recommendations of how phytochemical diversity can be measured with different types of datasets, and what tools are suitable to use.

An overview of our recommendations for measuring phytochemical diversity are presented in Figure 2. In general, we believe that although diversity should be measured with an appropriate index, it should also be deconstructed into separate measures of each component. Which components this includes depends on the dataset. The dataset may consist of presence/absence data, or quantitative data with the absolute/relative abundance of compounds. Setting aside disparity for now, in the former case the only measure of phytochemical diversity is the richness component. In the latter case, also the evenness of the compounds can be considered, if their relative abundances have been adequately quantified. With quantitative data, the chemical diversity may be quantified as Hill diversity. The diversity order parameter (*q*) can then be shifted to control the sensitivity of the measure to the relative abundance of compounds. Although any value of *q* > 0 can be used to include evenness as a component, we recommend using *q* = 1 by default, as compounds are then weighted in proportion to their abundances. Using a diversity index has the advantage of summarizing the diversity into a single number, but because it may conceal independent variation in its components, we recommend also quantifying richness and evenness separately, and examining variation in or effects of each of these components.

**Figure 2.**
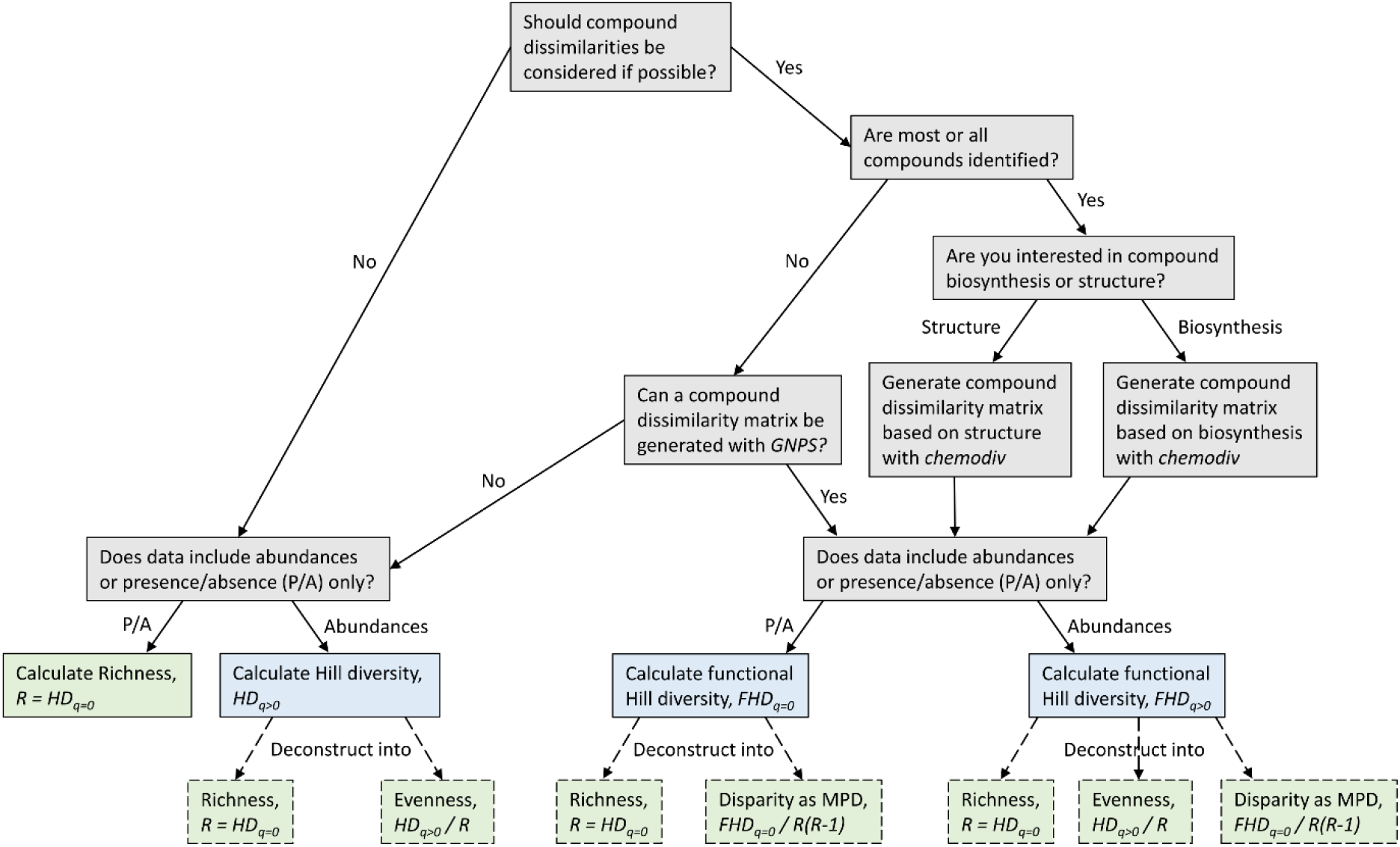
Decision tree for ways of quantifying phytochemical diversity. Choice of measure depends on if compound dissimilarity for the measured phytochemicals is to be used, and if abundance data, or only presence/absence data, is available. Grey boxes mark steps in the decision tree. Blue boxes represent measures of diversity, in the Hill diversity framework. Dashed green boxes mark components that measures of (functional) Hill diversity can be deconstructed into. Abbreviations: R: compound richness; HD_q=0_: Hill diversity at *q = 0*; HD_q>0_: Hill diversity at *q > 0*; FHD_q=0_: functional Hill diversity at *q = 0*; FHD_q>0_: functional Hill diversity at *q > 0*; MPD: mean pairwise dissimilarity. For measuring HD or FHD at *q* > 0, we recommend setting *q* = 1 by default. All steps in the decision tree (except generating a compound dissimilarity matrix with GNPS), can be done with the *chemodiv* R package.

In line with increasing evidence for the importance of compound structure for function, we think that the disparity of compounds (compound dissimilarity) should be considered if possible. This can be done in multiple ways. For datasets where most or all compounds remain unidentified, compound dissimilarities can be calculated as cosine scores based directly on mass spectra, by use of the GNPS ecosystem (Wang et al. 2016, Aksenov et al. 2021). For datasets where most or all compounds have been identified, we have previously developed the *chemodiv* R package (Petrén et al. 2023), which can be used to calculate compound dissimilarities based on either the structure of compounds, or their biosynthetic classification. Once a dissimilarity matrix of the compounds has been created, we recommend calculating functional Hill diversity (with *q* = 1 by default) to generate an overall measure of phytochemical diversity. Thereafter, this may be deconstructed into the three components. Richness and evenness is calculated in the same way as before, while disparity may be quantified as the mean pairwise dissimilarity (MPD). The *chemodiv* package also provides functions for these calculations.

The recommendations above represents a comprehensive way to measure chemodiversity. However, depending on what types of compounds constitute the dataset and the context it has been collected in, some measures are likely to be more relevant and interesting to examine than others. Below, we discuss measures of chemodiversity in these contexts.

### Aspects of the functional role of phytochemicals

Phytochemicals are functionally important in a wide range of interactions between plants and their environment, and the importance of different aspects of this chemodiversity may vary extensively depending on what type of interactions or interactors are considered. Although it may be difficult to answer which type of diversity is important in which context, some predictions can be made. In general, specialist herbivores are often less affected by chemical defences compared to generalists (Hopkins et al. 2009, Mithöfer and Boland 2012). This may be true also for phytochemical diversity (Dyer 2018). Several studies have found that phytochemical diversity can have a stronger negative effect on generalist than specialist herbivores (Agrawal 2000, Volf et al. 2018, Li et al. 2020, Bakhtiari et al. 2021, Kozel et al. 2022). It is important to note that such comparisons are made from the perspective of the plant(s) to which the specialists are specialised. Further, results are not always clear and may depend on both individual and community level diversity (Glassmire et al. 2020, Massad et al. 2022, Salazar and Marquis 2022). Phytochemical diversity might also be more predictive of function in comparisons within rather than between plant species (Schuldt et al. 2012). Within species, there may be limited variation in what compounds are found in different individuals, and function might thereby be more dependent on diversity. Across species, there may be more variation in what compounds are found in different species, which may be of greater importance for function than the diversity itself. It should also be mentioned that phytochemical variation on the level of whole plant communities rather than individual plants may also be ecologically important, where e.g. community level phytochemical similarity, diversity and uniqueness may affect herbivore diversity, levels of herbivory, plant survival and ecosystem functioning (Lavandero et al. 2009, Massad et al. 2017, Schuldt et al. 2018, Salazar and Marquis 2022) (Box 2).

From the perspective of the plant, phytochemical diversity might be more predictive of function under natural conditions when plants experience a multitude of interactions, compared to for single interactions. The value of phytochemical diversity for single interactions rests on the assumption that diversity *per se* is important for function, with synergistic effects of combinations of compounds. While such synergy might be common (Richards et al. 2016), there are also examples where synergistic effects are lacking (Liu et al. 2017, Whitehead et al. 2021), or where instead antagonistic effects are found (Whitehead and Bowers 2014, Heiling et al. 2022). When instead considering multiple interactions, phytochemical diversity can be predictive of function without direct synergistic effects, if different compounds are efficient against different interacting organisms (Berenbaum and Zangerl 1996, Iason et al. 2011). On the whole, we believe that phytochemical diversity is one important aspect of the phenotype, but also other aspects of the phytochemical phenotype are clearly key to function in many cases (Meier and Bowman 2008, Torres-Gurrola et al. 2011, Junker 2016). Chemodiversity should be measured as completely as possible and be deconstructed into its components, to exhaustively test for effects on function and its importance for shaping ecological interactions.

### Volatile phytochemicals as signals and cues

An important distinction regarding the function of phytochemicals in interactions is that compounds may function as toxins, or as information (Raguso et al. 2015, Kessler and Kalske 2018), which for insect herbivores affect their performance and preference, respectively. Although such a classification is a simplification, this distinction is important because the role of phytochemical diversity might differ for compounds that when consumed have direct negative physiological effects on consumers (Wari et al. 2021), compared to the most often volatile compounds acting as cues or signals providing information to herbivores and pollinators (Wilson et al. 2015). Among studies testing for functional effects of phytochemical diversity, only a few have done it in an information context (Table S1), examining how variation in both individual and community level volatile diversity can deter herbivores (e.g. Doyle 2009, Zu et al. 2020, 2022, Galmán et al. 2022) or attract parasitoids of herbivores (Wan et al. 2017). In the case where phytochemical compounds act as floral scent to attract pollinators, some studies have examined variation in richness between e.g. different populations (Zeng et al. 2022, Eisen et al. 2022) or compared it for plants pollinated by different groups of pollinators (Farré-Armengol et al. 2020). However, to our knowledge, no study has empirically tested for a direct effect of phytochemical diversity on pollinator attraction.

It is possible that some components of the chemodiversity are more or less relevant to the functional aspects of the phytochemical phenotype when this constitutes information rather than toxins. Phytochemical richness might be more relevant than evenness, as also low abundance compounds can be highly attracting or repelling to insects (Clavijo McCormick et al. 2014). The disparity may also be an important component. Because different classes of compounds may, on a general level, have partly different functions in plant-insect interactions (Schiestl 2010, Junker and Blüthgen 2010, Kantsa et al. 2019), a set of biosynthetically diverse compounds could be more multifunctional than a set of compounds produced in the same biosynthetic pathway. In general, we need a better understanding about what aspects of the phenotype, including individual compounds (Zhou et al. 2017), specific ratios/compositions (Wright et al. 2005, Orlando et al. 2022), modules of compounds (Junker et al. 2018), or the diversity of compounds, are functioning as information cues or signals sent out by plants. Related to this, as communication via volatiles is susceptible to noise (Wilson et al. 2015), it should also be examined whether phytochemical diversity *per se*, through synergies between compounds, is similarly important to function when phytochemicals function as information compared to toxins.

### Evolutionary patterns

From an ecological perspective, phytochemicals affect the organisms interacting with plants. From an evolutionary perspective, the interacting organisms are agents of selection, affecting what phytochemicals plants produce. By examining macroevolutionary patterns and microevolutionary processes, we can learn how natural selection and other forces of evolution generate chemodiversity on different levels.

Assuming a coevolutionary escape-and-radiate process between plants and herbivores (Ehrlich and Raven 1964), the diversity of phytochemicals should increase over time as new plant species evolve (Speed et al. 2015). A few studies have found evidence of this. Becerra et al. (2009) found that the phytochemical diversity in *Bursera* species, calculated based on both the number and types of compounds, escalated over macroevolutionary timescales. Similarly, Volf et al. (2018) found an increase in alkaloid diversity over time among *Ficus* species, and Defossez et al. (2021) discovered an increase in the richness of molecular families over time in a set of 416 vascular plant species. In contrast, Cacho et al. (2015) found an evolutionary decline in glucosinolate diversity in *Streptanthus* plants, suggesting potential trade-offs with other kinds of defence (Agrawal and Fishbein 2008). These studies suggest that phytochemical diversity may often, but not always, increase over evolutionary time. This can include new compounds produced in existing biosynthetic pathways, or, potentially more efficient but less common, compounds produced in new biosynthetic pathways (Becerra et al. 2009). Overall, phylogenetic patterns are often complex and might differ for different aspects of the phytochemical phenotype, or different classes of compounds (Courtois et al. 2016, Züst et al. 2020, Zhang et al. 2021, Forrister et al. 2022). Therefore, the disparity component of chemodiversity, based on biosynthetic classifications, should be taken into account in studies examining macroevolutionary patterns of phytochemicals.

Few studies investigate patterns and processes surrounding chemodiversity on a microevolutionary scale. The two examples that have investigated associations between (neutral) genetic diversity and phytochemical diversity have found different results (Bravo-Monzón et al. 2018, Pais et al. 2018). Direct tests for phenotypic selection on phytochemical diversity are lacking. While multiple studies have examined selection on individual compounds, principal components or total abundances of e.g. floral scent or herbivore defence compounds (Johnson et al. 2009, Chapurlat et al. 2019, Joffard et al. 2020), we are aware of no studies that have measured phenotypic selection on phytochemical diversity directly. There are multiple examples of associations between phytochemical diversity and various measures of plant performance, such as levels of herbivory. However, these do not take into account the potential metabolic or ecological costs for plants to produce a diverse set of compounds, which could outweigh benefits (Cipollini et al. 2017). Therefore, direct estimates of selection are required to investigate the potential for phenotypic selection to act on a composite trait such as phytochemical diversity (*c.f.* Opedal et al. 2022). Doing so, we may gain a better understanding of how different aspects of the phytochemical phenotype function ecologically, come under selection, and evolve over time.

### Phenotypic plasticity

Phytochemical diversity is not only genetically determined, but may also be affected by the plant’s surrounding environment. Phenotypic plasticity of phytochemicals, and its adaptive value, has been examined in a broad set of contexts, including inducibility by herbivores, temporal variation and variation due to abiotic factors (Majetic et al. 2009, Metlen et al. 2009). Any phenotypic change is likely to have an effect on chemodiversity, but direct studies on this are rare, with only a handful of examples. Agrawal (2000) found an increase in glucosinolate richness following herbivory in *Lepidium virginicum*, which was associated with a decreased performance in a generalist, but not a specialist, herbivore. In contrast, Li et al. (2020) and Bai et al. (2022) found that herbivory in *Nicotiana attenuata* had no or a negative effect on the diversity of leaf metabolomes. Instead, induced changes may act to increase metabolomic specialization, if there is an increased production of certain groups of compounds that increase plant resistance. Changes to the abiotic environment could also affect levels of chemodiversity (Ramos et al. 2021), although Tewes and Müller (2018) found no effect of fertilization on glucosinolate diversity in *Bunias orientalis*.

Overall, although in plants chemical traits may be more plastic, at least over short temporal scales, than morphological traits (Walker et al. 2022), we still know comparatively little about what dimensions of the phytochemical phenotype are most plastic. If plasticity primarily involves changes in the relative abundances of compounds, this will affect phytochemical evenness. If it involves production of other sets of compounds, the richness or disparity components may instead change. More studies are needed to examine the plasticity of different aspects of the phenotype, investigate the adaptive value of such plasticity, and better distinguish between genetic and environmental causes of variation.

### Unanswered questions and future research directions

A number of aspects of phytochemical diversity and surrounding topics have various central questions that remain to be more closely investigated. First, as mentioned in previous sections, phytochemical diversity is only one aspect of a multivariate phytochemical phenotype. To what extent the diversity, in contrast to e.g. specific compounds, compositions of compounds, classes of compounds or the total abundance of compounds, is mechanistically important for, or simply predictive of, function will be a central question to answer (Yarnes et al. 2006, Marion et al. 2015, Oduor 2022). Crucial to this will be to better understand the links between molecular structure and function, further examine the role of synergistic effects, and test if structurally diverse sets of compounds also have broader function or more potent effects (Berenbaum and Zangerl 1996, Liu and Zhao 2016, Philbin et al. 2022). Understanding this, in turn, requires an increased understanding on how phytochemicals function mechanistically on a physiological and molecular level (Mithöfer and Boland 2012, Wari et al. 2021).

A second important aspect is diversity specifically. As diversity is a composite measure, we have argued for first measuring it, and then deconstructing it into its components of richness, evenness and disparity. Many alternative and extended ways of quantifying diversity exist, that may also provide relevant measures of the phytochemical phenotype (Mouchet et al. 2010, Chao et al. 2019, Magneville et al. 2021), although more complex measures may be more difficult to interpret. Additionally, different types/components of diversity may be mathematically correlated, and if comparing multiple measures care should be taken to not mistake mathematical associations for biological ones (Loiseau and Gaertner 2015). Other quantities related to diversity, such as specialization and dominance, may also be important (Berger and Parker 1970, Martínez and Reyes-Valdés 2008), as may be measures of intramolecular complexity, which are not quantified by measures of compound dissimilarity (Richards et al. 2015, Méndez-Lucio and Medina-Franco 2017, Philbin et al. 2022). In addition, although we have focused on variation in phytochemical diversity at the level of individual plants, also β-diversity or diversity at the population or community is relevant for ecological function (Wetzel and Whitehead 2020, Glassmire et al. 2020, Robinson et al. 2022). Studies that simultaneously quantify multiple types/components of diversity are needed to examine which of these quantities are most relevant in different contexts.

Third, studies on phytochemical diversity remain somewhat limited in scope in regards to what species are studied and which experimental methods are used. The vast majority of studies have been done on flowering plants, with only a few examples including gymnosperms or mosses (Iason et al. 2005, Peters et al. 2018, 2019, 2021, Feng et al. 2021). Regarding how studies are designed, a large part of them are observational, comparing chemodiversity across different groups and/or associating this with ecological function. This is most often done on the level of individual plants, and further examinations at both smaller and larger spatial scales, from within plants to between communities, are needed. Experimental studies manipulating levels of diversity are rare (Whitehead et al. 2021, Fernandez-Conradi et al. 2022, Salazar and Marquis 2022), but useful for disentangling what components of diversity are most relevant for function. Additionally, most research on the effects of phytochemical diversity examine the effect of this diversity in leaves on the performance of insect herbivores or levels of herbivory. Studies including other plant parts such as roots, fruit and seeds, and other types of interactions such as those involving fungi, bacteria, or in an information context where phytochemicals acts as signals or cues, are more rare (Lindig-Cisneros et al. 1997, 2002, Doyle 2009, Pais et al. 2018, De-la-Cruz-Chacón et al. 2019, Zu et al. 2020, Whitehead et al. 2021, Feng et al. 2021). Additionally, recent technical advances in plant metabolomics enable the use of new analytical methods (Uthe et al. 2021), which will be valuable for future insights. Finally, although most research so far has aimed to answer fundamental ecological questions, a few studies have been done in a more applied context, examining phytochemical diversity for crop species, and its potential importance for protection against pest insects (Whitehead and Poveda 2019, Espinosa-García et al. 2021, Robinson et al. 2022, Bernal et al. 2023). Further research on this topic could increase our understanding of how considerations of chemodiversity could be utilized in agroecosystems (Silva et al. 2018, Espinosa-García 2022).

## Conclusions

More than 60 years have passed since Fraenkel’s (1959) seminal paper on the raison d’être of phytochemical compounds. Today, studying the mechanisms by which these phytochemicals function, in order to explain patterns or effects observed in nature, is a central topic in chemical ecology (Raguso et al. 2015). However, we still have a limited knowledge of how these compounds, alone or in mixtures, function in different types of interactions between plants and their environment. As evident by the review of the literature, considering mixtures of phytochemicals as a complex phenotype, where aspects of its multivariate nature can be summarized into measures of diversity, can be a fruitful way of increasing our understanding of phytochemical function. To achieve a more nuanced and complete understanding of this, we have made recommendations of how to measure phytochemical diversity in different contexts, which will allow researchers to better study the relevant parts of phytochemical variation. In this way, we hope to contribute to an increased understanding of the functional importance of the diversity of phytochemicals produced by plants.

Box 1 Glossary of central terms

*Phytochemicals* – also referred to as plant secondary compounds or specialized (secondary) metabolites. These are compounds produced by plants, which function predominantly in (a)biotic interactions between plants and their environment, rather than being part of primary metabolic functions.

*Phytochemical phenotype* – the combined set of phytochemical compounds found in or emitted by (part of) a plant, with each compound representing a “trait” making up the complete multivariate phenotype.

*Phytochemical diversity* (*Chemodiversity*) – the diversity of a set of phytochemical compounds, which (if the set is from an individual plant) represents an aspect of the phytochemical phenotype. Diversity itself is a multifaceted concept, and in this study, we regard diversity as some combination of richness, evenness and disparity.

*Phytochemical richness* – a measure of the number of unique compounds in a sample.

*Phytochemical evenness* – a measure of the equitability of the relative abundances of compounds in a sample. Evenness is high when all compounds have equal abundances, and low when one compound has a high abundance and others have a low abundance.

*Phytochemical disparity –* a measure of how dissimilar the compounds in a sample are. We refer to this as compound dissimilarity, and this can be quantified based on e.g. the biosynthetic classification or structural properties of the compounds. A pair of compounds have a pairwise dissimilarity, and all the pairwise dissimilarities for a set of compounds can be used to construct a compound dissimilarity matrix.

*Diversity index* – a quantitative measure of diversity. There are many different diversity indices, which in different ways quantify diversity as a function of richness, evenness and/or disparity.

*Functional diversity –* although a term with varying meanings, we use the following definition: for species diversity, functional diversity describes the diversity of functional traits of species in a community. For phytochemical diversity, it describes the diversity of (functional) properties of compounds in a sample. In practice, here we regard a diversity index to quantify functional diversity if it includes a disparity component in the measure, where disparity is based on the structure or biosynthesis of compounds. Assuming a link between dissimilarity in structure/biosynthesis and dissimilarity in function, such a measure then quantifies the diversity of functions of the compounds in a sample.

*Hill diversity* - also called Hill numbers or effective number of species, these are a set of diversity indices, expressed in units of effective numbers, that have multiple advantages over traditional diversity indices such as Shannon’s diversity or Simpson diversity. With the use of (functional) Hill diversity, each of the three components of diversity can be quantified separately and combined.

Box 2 Similarities of, and links between, species diversity and phytochemical diversity

In this paper, we review the application of the concept of diversity, most often used to measure the diversity of species, to instead measure the diversity of phytochemicals. There are several conceptual similarities between these applications that deserves to be mentioned.

The diversity of species is of interest both as a measure in itself, and, importantly, due to the effect of biodiversity on ecosystem functioning. A large body of literature has demonstrated positive effects of diversity, mostly in plants, on a wide range of ecosystem functions in a wide range of systems (Loreau et al. 2001, van der Plas 2019). These studies indicate that different components of diversity, including the richness, evenness and disparity, can be important for ecosystem function in different contexts (Tilman et al. 2014), although functional diversity is often found to be more important for function than species diversity (Tilman et al. 1997, van der Plas 2019). Mechanistically, diversity might have positive effects on ecosystem function in different ways, including “complementarity effects” where the functioning of individual species is higher when grown in communities rather than monocultures, and “selection effects”, where communities with high species richness are more likely to include (high abundances of) highly functioning species (Loreau and Hector 2001). Additionally, the functional composition of species is also often a major factor affecting ecosystem function (van der Plas 2019).

The diversity of phytochemicals, and its effects on plant function, works in analogous ways. We have demonstrated (see section “Systematic literature review”) how different components of diversity can influence plant function, and argued that measures of functional diversity, that include a disparity component, might be more predictive of function than measures based on only richness and/or evenness. Mechanistically, there are connections as well. The “synergy hypothesis” (synergistic effects between compounds, see section “Phytochemical richness”) is comparable to the “complementarity effects”. The “screening hypothesis” (only few compounds are functional) is related to the “selection effects”. The “interaction diversity hypothesis” (different compounds are efficient against different interacting organisms), is related to the concept of ecosystem multifunctionality, where different species affect different ecosystem processes (Hector and Bagchi 2007). In the same way, in addition to the diversity, also the composition of compounds is likely often important for function as well. The similarities between effects of species diversity and phytochemical diversity demonstrate an interesting generality of diversity-function relationships.

While we have considered species and phytochemical diversity separately above, the concepts can also be linked to each other. Phytochemicals can, similar to morphological traits, be regarded as (functional) traits (Müller and Junker 2022, Walker et al. 2022). For measures of phytochemical diversity, these traits collectively make up the phenotype, which can be quantified as a measure of its diversity. In contrast, for calculations of the functional diversity of species in a community, these traits are instead included in calculations of the functional (chemo)diversity of species. Studies utilizing phytochemicals as functional traits, or simply quantifying phytochemical diversity on a community rather than individual plant level, have found effects of this diversity on plant functions such herbivore resistance, which itself can be regarded as an ecosystem function (Salazar et al. 2016, Schuldt et al. 2018, Fernandez-Conradi et al. 2022, Ristok et al. 2023). In this way, the diversity of the phytochemicals that plants produce constitutes a part of the mechanistic link between plant diversity and ecosystem function.

## Author contributions

RRJ and HP conceived of the study. All authors contributed ideas and perspectives on the topics of the paper. HP performed the systematic literature review and wrote the first draft with contributions from RRJ. All authors reviewed and approved the submitted version of the manuscript.

## Acknowledgements

This study was done within the Research Unit FOR3000, funded by the German Research Foundation (Deutsche Forschungsgemeinschaft, JU 2856/5-1).

## Conflict of interest

The authors declare no conflict of interest.

## Notes

### Competing Interest Statement

The authors have declared no competing interest.

