## Supplementary material for "Understanding the phytochemical diversity of plants: Quantification, variation and ecological function": Table S1

**Table S1.** List of studies on phytochemical diversity found in the systematic literature review, including study system, type of phytochemicals measured, analytic method, type of diversity or component thereof measured, and if variation in, or an effect of, diversity was found or not.

| General information | | | | Variation | | Effects | | | | Ref. |
|---|---|---|---|---|---|---|---|---|---|---|
| **Study system** | **Phytochemicals measured** | **Analytic method** | **Measures used** | **Group variation** | **No group variation** | **Positive effect** | **Negative effect** | **No effect** | **Complex effect** | |
| Artificial diet | Various compounds | - | Richness | | | Y | | Y | | 1 |
| *Lepidium virginicum* | Glucosinolates | GC-FID | Richness | Y | | Y | | | | 2 |
| *Nicotiana attenuata* | Leaf compounds | LC-MSMS | Shannon | Y | | | | | | 3 |
| *Cardamine* | Leaf glucosinolates | LC-MS | Richness, evenness, Shannon, Rao's Q, FRic, FDiv | Y | | Y | Y | Y | | 4 |
| Fabaceae, Senecioneae | Various compounds | - | Richness, Shannon | Y | | | | | | 5 |
| *Bursera* | Leaf VOCs | GC-MS | Shannon, complexity index | Y | | | | | | 6 |
| *Zea mays* | Root VOCs | GC-MS | Evenness, Simpson | Y | | | | | Y | 7 |
| *Mikania micrantha* | Leaf terpene VOCs | GC-MS | Shannon | Y | | | | | | 8 |
| *Mikania micrantha* | Leaf terpene VOCs | GC-MS | Shannon | Y | | | | | | 9 |
| *Streptanthus* | Leaf glucosinolates | HPLC | Richness, Shannon, complexity index | Y | | | | | | 10 |
| Artificial diet | Various compounds | - | Richness | | | Y | | | | 11 |
| *Piper amalago* | Leaf compounds | LC-MS, NMR | Hill-Shannon | | | Y | | | | 12 |
| 55 tree species | Leaf VOCs | GC-MS | Richness | Y | | | | | | 13 |
| 202 tree species | Leaf terpene VOCs | GC-MS | Richness | Y | | | | | | 14 |
| *Annona purpurea* | Leaf, stem, root alkaloids | GC-MSMS | Margalef | | | Y | | | | 15 |
| 416 plant species | Leaf compounds | LC-MSMS | Richness | Y | | Y | | | | 16 |
| *Gossypium hirsutum* | Leaf VOCs | GC-MS | Shannon (joint entropy) | | | | | | Y | 17 |

| General information | | | | Variation | | Effects | | | | Ref. |
|---|---|---|---|---|---|---|---|---|---|---|
| **Study system** | **Phytochemicals measured** | **Analytic method** | **Measures used** | **Group variation** | **No group variation** | **Positive effect** | **Negative effect** | **No effect** | **Complex effect** | |
| *Erodium cicutarium* | Leaf, blossom, fruit terpenes | GC-MS | Shannon | Y | | | | | | 18 |
| *Tanacetum vulgare* | Floral VOCs, pollen compounds | GC-MS | Richness, Shannon | Y | | | | | | 19 |
| *Oenothera cespitosa* | Floral VOCs | GC-MS | Richness | Y | | | | | | 20 |
| *Nicotiana* | Leaf, root, calyx compounds | LC-MSMS | Richness, Shannon | Y | | | | | | 21 |
| *Persea americana* | Leaf compounds | GC-MS | Evenness, Shannon, Simpson | | | Y | | Y | | 22 |
| 305 plant species | Floral VOCs | - | Richness | Y | | | | | | 23 |
| *Juniperus rigida* | Needle compounds | GC-MS | Evenness, Shannon, Simpson | Y | | Y | | Y | | 24 |
| 37 plant species | Leaf compounds | LC-MSMS | Richness | | | Y | Y | Y | | 25 |
| *Inga* | Leaf compounds | LC-MSMS | Functional Hill diversity | Y | | | | | | 26 |
| *Quercus ilex* | Leaf VOCs, phenolics | HPLC, GC-MS | Shannon | Y | | | | Y | | 27 |
| *Piper kelleyi* | Leaf compounds | HPLC | Hill-Shannon | Y | | | | | Y | 28 |
| *Piper kelleyi* | Leaf compounds | LC-MS, NMR | Richness, Simpson | Y | | Y | | | | 29 |
| *Solanum Pennellii* | Leaf compounds | LC-MS | Simpson | | | Y | | | | 30 |
| *Medicago sativa* | Leaf compounds | HPLC | Hill-Shannon | | Y | | | Y | | 31 |
| *Medicago sativa* | Leaf compounds | LC-MS | Hill-Shannon | | | | | | Y | 32 |
| *Pinus sylvestris* | Needle terpenes | GC-MS | Shannon, Simpson | | | | | | Y | 33 |
| Apiaceae | Classes of compounds | - | Richness | | | | | Y | | 34 |
| *Lupinus polyphyllus* | Leaf alkaloids | HPLC | Richness | | Y | | | | | 35 |
| *Salix* | Leaf compounds | LC-MSMS | Richness | Y | | Y | | | | 36 |
| *Nothofagus* | Leaf compounds | TLC | Richness | | | | | Y | | 37 |

| General information | | | | Variation | | Effects | | | | Ref. |
|---|---|---|---|---|---|---|---|---|---|---|
| **Study system** | **Phytochemicals measured** | **Analytic method** | **Measures used** | **Group variation** | **No group variation** | **Positive effect** | **Negative effect** | **No effect** | **Complex effect** | |
| *Nicotiana attenuata* | Compounds (different tissues) | LC-MSMS | Shannon | Y | | | | | | 38 |
| *Nicotiana attenuata* | Leaf compounds | LC-MSMS | Shannon | Y | | | | | | 39 |
| *Phaseolus* | Leaf compounds | HPLC | Shannon | | | Y | | | | 40 |
| *Phaseolus vulgaris* | Leaf compounds | HPLC | Shannon | | | Y | | | | 41 |
| 12 Asteraceae species | Leaf compounds | LC-MS | Richness | Y | | | | | Y | 42 |
| *Piper* | Leaf compounds | NMR | Simpson | | | | | | Y | 43 |
| *Erysimum cheiranthoides* | Leaf cardenolides | LC-MS | Evenness | Y | | | | | | 44 |
| *Plantago lanceolata* | Leaf compounds | LC-MS | Richness, evenness, Shannon, Simpson | | | | | | Y | 45 |
| *Brassica nigra* | Leaf glucosinolates | HPLC | Shannon | Y | | | | | Y | 46 |
| *Cecropia* | Leaf compounds | HPLC | Richness, Shannon | Y | | | | | | 47 |
| *Cornus florida* | Leaf compounds | LC-MS | Richness, evenness, Shannon, Simpson, Berger-Parker | Y | | | Y | | | 48 |
| 8 trees species | Leaf compounds | LC-MS | Richness, Shannon | Y | | | | | | 49 |
| 9 bryophyte species | Moss compounds | LC-MS | Shannon | Y | | | | | | 50 |
| 9 bryophyte species | Moss compounds | LC-MSMS | Richness, evenness, Shannon | Y | | | | | | 51 |
| 10 bryophyte species | Moss compounds | LC-MSMS | Richness, evenness, Shannon | Y | | | | | | 52 |
| *Ceanothus velutinus* | Leaf compounds | LC-MS | Hill-Shannon | Y | | | | | Y | 53 |
| *Piper* | Leaf compounds | LC-MS, NMR | Hill-Shannon | Y | | Y | | Y | Y | 54 |
| *Piper* | Leaf compounds | GC-MS | Shannon | Y | | | | | | 55 |
| *Apocynum* | Root cardenolides | HPLC | Shannon | Y | | | | | Y | 56 |

| General information | | | | Variation | | Effects | | | | Ref. |
|---|---|---|---|---|---|---|---|---|---|---|
| **Study system** | **Phytochemicals measured** | **Analytic method** | **Measures used** | **Group variation** | **No group variation** | **Positive effect** | **Negative effect** | **No effect** | **Complex effect** | |
| *Asclepias* | Leaf and root cardenolides | HPLC | Shannon | Y | | | | | | 57 |
| *Piper* | Leaf compounds | NMR | Simpson | | | Y | | | | 58 |
| 8 grassland species | Leaf compounds | LC-MS | Richness, Hill-Shannon | Y | | Y | Y | | | 59 |
| *Medicago sativa* | Leaf saponins | LC-MSMS | Richness, Shannon, Simpson | Y | | | | | | 60 |
| *Piper* | Leaf compounds | GC-MS | Rao's Q (community level) | | | Y | | | | 61 |
| 31 Burseraceae species | Leaf compounds | GC-MS, LC-MS | Richness | Y | | Y | | | | 62 |
| *Piper* | Leaf compounds | - | Rao's Q (community level) | | | Y | | Y | | 63 |
| Various plant species | Floral VOCs | - | Richness, Shannon | | | | | | Y | 64 |
| *Piper* | Leaf and fruit compounds | LC-MSMS | Richness | Y | | | | | | 65 |
| 21 tree/shrub species | Leaf phenolics and tannins | HPLC | Shannon | | | | | Y | | 66 |
| *Piper* | Leaf compounds | NMR | Simpson | | | | | | Y | 67 |
| *Asclepias* | Leaf cardenolides | HPLC | Shannon | | | | | | Y | 68 |
| *Bunias orientalis* | Leaf glucosinolates | LC-MS | Shannon | | | Y | | | | 69 |
| *Bunias orientalis* | Leaf glucosinolates | LC-MS | Shannon | Y | Y | | | | | 70 |
| *Persea americana* | Leaf compounds | GC-FID | Shannon | | | | | Y | | 71 |
| *Ficus* | Leaf compounds | LC-MS(MS) | Shannon | Y | | | | | Y | 72 |
| *Ficus* | Leaf compounds | LC-MS(MS) | Shannon | Y | | | | | Y | 73 |
| *Salix* | Leaf compounds | LS-MSMS | MPD | Y | | | | | | 74 |
| *Glycine max* | Plant volatiles | GC-MS | Shannon, Simpson, Brillouin, McIntosh | Y | | Y | | | | 75 |
| 4 tree species | Leaf and root compounds | LC-MS | Richness, Shannon | Y | | | | | | 76 |

| General information | | | | Variation | | Effects | | | | Ref. |
|---|---|---|---|---|---|---|---|---|---|---|
| **Study system** | **Phytochemicals measured** | **Analytic method** | **Measures used** | **Group variation** | **No group variation** | **Positive effect** | **Negative effect** | **No effect** | **Complex effect** | |
| *Malus* | Fruit phenolics | HPLC | Shannon | | | Y | | | | 77 |
| *Piper reticulatum* | Leaf, root, flower, fruit, seed amides | GC-MS | Richness | Y | | | | | | 78 |
| Artificial diet | Phenolics | - | Richness, evenness, MPD | | | Y | | Y | | 79 |
| *Tanacetum vulgare* | Leaf terpenes | GC-MS | Shannon, evenness (of chemotypes) | Y | | | | | | 80 |
| *Tanacetum vulgare* | Leaf compounds | GC-MS | Shannon | Y | Y | | | | | 81 |
| *Ficus* | Fig VOCs | - | Shannon (conditional entropy) | | | | | | Y | 82 |
| *Primula Oreodoxa* | Floral VOCs | GC-MS | Richness | Y | | | | | | 83 |
| 315 tree species | Root compounds | LC-MSMS | Shannon | Y | | | | | Y | 84 |
| *Tanacetum vulgare* | Leaf terpenes | GC-MS | Shannon (individual and community level) | | | Y | Y | Y | | 85 |
| 20 plant species | Leaf VOCs | GC-MS | Shannon (conditional entropy) | | | | | | Y | 86 |
| *Erysimum* | Leaf glucosinolates, cardenolides | GC-MS | Richness | Y | | | | | | 87 |

*Study system* describes the system in which the study was performed. *Phytochemicals measured* describes what kind of tissue samples were taken from, and what kind of compounds were measured (if applicable; this includes biosynthetic class and if compounds were Volatile Organic Compounds (VOCs)). *Analytic method* describes what kind of analytic method compounds were analysed with. *Measures used* describes which diversity indices, or components of diversity, were measured (MPD: mean pairwise dissimilarity, Hill-Shannon: Hill diversity at q = 1, which corresponds to the exponential of Shannon's diversity). *Group variation* indicates that variation was found between groups, e.g. populations, species, treatments and tissues (*Y* indicates Yes). *No group variation* indicates that no variation was found between groups, e.g. populations, species, treatments and tissues, when it was tested for. *Positive effect*, *Negative effect* and *No effect* indicates whether there was direct evidence of positive, negative or no effect, respectively, of an increased phytochemical diversity on some aspect (e.g. herbivore resistance) of plant performance. *Complex effect* indicates that (according to our best judgements) effects were variable, complex or difficult to interpret, with uncertain effects on plant performance, or effects that were difficult to relate to plant performance. *Ref.* indicates the reference for the study. If a study tested for variation, there is a *Y* for *Group variation* and/or *No group variation*. If both these cells are empty, this indicates variation was not tested for/not applicable. If a study tested for an effect, there is a *Y* for *Positive effect*, *Negative effect*, *No effect* and/or C*omplex effect*. If all these four cells are empty, this indicates an effect was not tested for/not applicable.
